# Single Cell MALDI-MSI Analysis of Lipids and Proteins within a Replicative Senescence Fibroblast Model

**DOI:** 10.1101/2024.03.13.584140

**Authors:** Emily R. Sekera, Lorena Rosas, Joseph H. Holbrook, Quetzalli D. Angeles-Lopez, Timur Khaliullin, Mauricio Rojas, Ana L. Mora, Amanda B. Hummon

## Abstract

In this study, we evaluate lipids and select proteins in human lung fibroblasts (hLFs) to interrogate changes occurring due to aging and senescence. To study single cell populations, a comparison of cells adhered onto slides using poly-D-lysine versus centrifugal force deposition was first analyzed to determine whether specific alterations were observed between preparations. The poly-D-lysine approach was than utilized to interrogate the lipidome of the cell populations and further evaluate potential applications of the MALDI-immunohistochemistry (IHC) platform for single-cell level analyses. Two protein markers of senescence, vimentin and p21, were both observed within the fibroblast populations and quantified. Lipidomic analysis of the fibroblasts found twelve lipids significantly altered because of replicative senescence, including fatty acids, such as stearic acid, and ceramide phosphoethanolamine species (CerPE). Similar to previous reports, alterations were detected in putative fatty acid building blocks, ceramides, among other lipid species. Altogether, our results reveal the ability to detect lipids implicated in senescence and show alterations to protein expression between normal and senescent fibroblast populations, including differences between young and aged cells. This report is the first time that the MALDI-IHC system has been utilized at a single-cell level to analyze both protein expression and lipid profiles in cultured cells, with a particular focus on changes associated with aging and senescence.

## INTRODUCTION

Aging is a complex process occurring due to changes at the biological, biophysical, and biochemical levels, in part due to the accumulation of senescent cells. Senescence is a cell fate decision, characterized by cell cycle arrest, resistance to apoptosis and the senescence-associate secretory phenotype (SASP). Although senescence has beneficial physiological effects during development, wound healing, and in cancer prevention; persistence of senescence contributes to the development of fibrotic and age-related diseases^1^. Several triggers of senescence have been described, including the gradual loss of telomeres due to successive cell division, a process known as replicative senescence^2^.

First described by Hayflick and Moorhead in the early 1960’s, their study of human fetal fibroblasts in culture noted that cells gradually lost their ability to proliferate upon continued subculturing^2^. This phenomenon reflects the finite life cycle of somatic cells. As cells undergo senescence, morphological and functional alterations can be observed, including the enlargement and flattening of cells and stable cell cycle arrest. To detect senescence within cells, a multi-faceted panel of markers is needed to encapsulate the variability between cell populations. Most commonly, the identification of senescent cells relies on the increased expression of markers such as senescence-associated β-galactosidase (SA-β-gal) and cell cycle arrest regulators such as p16^INK4A^, and p21^WAF1/Cip1^. These proteins are often used alongside markers for the SASP, which includes various secreted factors involved in inflammation and tissue remodeling^3^.

To understand cellular senescence comprehensively, it is essential to examine both lipid metabolism and traditional senescence markers. Senescent cells exhibit changes in lipid profiles, including the accumulation of specific lipid species.^1^ These changes can influence cellular stress responses and the expression of stress-related proteins like vimentin^4, 5^. This interplay between altered lipid metabolism and protein expression is key to understanding how senescent cells maintain and propagate their altered state.

The analyses of these markers *in vivo* rely heavily upon techniques such as immunostaining that can provide spatial context to the biological questions interrogated. Integrating single-cell proteomics and lipidomics can provide insight into mechanisms underlying senescence and disease. Rapidly developing technology within the field of mass spectrometry imaging (MSI) has advanced the technique to the routine interrogation of single cells.^6^ In 2022, Cuypers et al. demonstrated the potential for utilizing single cell lipidomic analyses using MSI to differentiate between epithelial cell types and further utilized the unique lipid signals to differentiate between cell lines within a tumor xenograft model.^7^ Similarly, Capolupo and coworkers determined that lipid signals in fibroblasts could be used to determine the transcriptional programming of dermal cells.^8^ While lipid analysis is rapidly evolving across a multitude of MSI platforms for single cell analyses, protein imaging has proven to be more challenging. To address these challenges, the utilization of a matrix-assisted laser desorption/ionization immunohistochemistry (MALDI-IHC) platform allows for the labelling of accessible antigens with photocleavable mass-tags (PC-MTs) to determine putative localizations of proteins similar to traditional IHC workflows.

MALDI-IHC has been successfully applied to a range of tissues and continues to advance towards multi-modal imaging applications. The technique was first reported by Yagnik and colleagues to interrogate formalin-fixed paraffin-embedded (FFPE) mouse brain, human tonsil, and breast cancer tissue specimens. The authors also showcased the utility of the platform to run consecutive analyses on fresh frozen tissues for lipid and protein analyses.^9^ Claes *et al*. further interrogated the utility of MALDI-IHC to be used in conjunction with untargeted *in situ* bottom-up proteomics within breast cancer tissue.^10^ Moving towards multi-modal imaging analyses, dual labeled PC-MTs with fluorophores were introduced to enable complementary validation within the same samples.^11^ Further method development interrogating the use of antibody-directed single cell workflows, including GeoMx and imaging mass cytometry, in conjunction with MALDI-MSI were discussed by Dunne and coworkers within breast and liver tissue samples.^12^ Herein we discuss the utilization of a MALDI-IHC platform for relative quantitation of targeted proteins paired with lipidomic analysis of cultured fibroblasts. We examine human lung fibroblasts with and without induced replicative senescence, as well as from young and old individuals, to determine changes at a single-cell level and gain further insight into cell-to-cell variation.

## EXPERIMENTAL

### Preparation and Data Collection of Primary Human Lung Fibroblasts

Primary human lung fibroblasts (hLFs) were isolated from healthy young donor and healthy old donor tissues from the Comprehensive Transplant Center (CTC) Human Tissue Biorepository at The Ohio State University with Total Transplant Care Protocol informed consent and research authorization form. A table of patient demographics can be found in **Figure 1A** and the experimental workflow is shown in **Figure 1B**. CTC operates in accordance with NCI and ISBER Best Practices for Repositories. Details on cell culture and treatments, including induction of replicative senescence can be found in the supplementary information. Confirmation of senescence in samples was completed using senescence-β-galactosidase staining; experimental conditions and results can be found in the supporting information (**Figure S1**).

**Figure 1:**
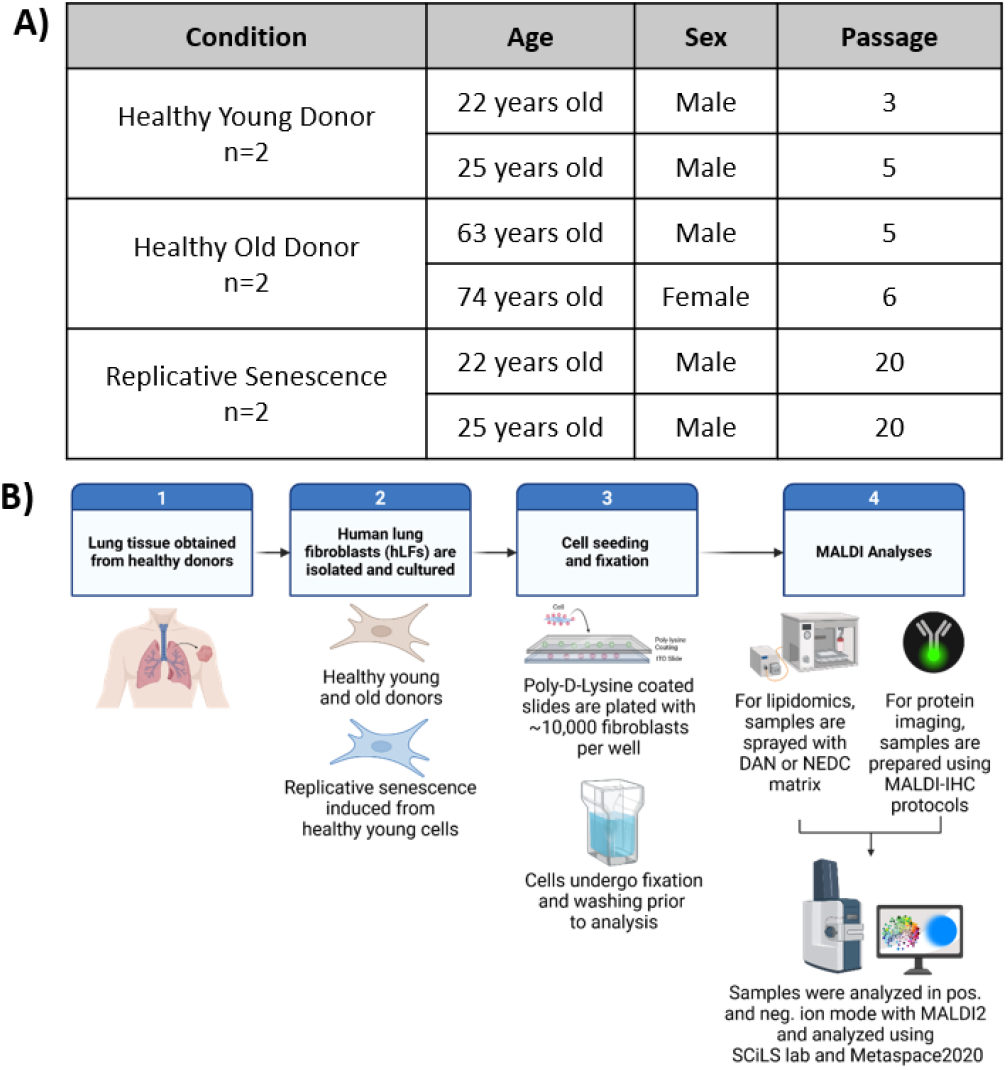
Donor demographics and experimental design. **A)** Description of condition, age, sex, and passage number for the hLFs utilized within the studies. **B)** Experimental overview for preparation of hLFs for lipidomic and proteomic analyses. Workflow created using BioRender.com.

### Fibroblast Preparation for Single Cell MSI

Fibroblasts were prepared for single cell MSI following either previous protocols^7^ or modified procedures. Cultured hLFs were lifted from slides using 0.05% trypsin, and either plated on poly-D-lysine slides for overnight adherence or deposited onto slides using a Cytospin™4 (Thermo Fisher Scientific, Carlsbad, CA) cytocentrifuge.

To aid cells in adhering to indium tin oxide (ITO) slides for MALDI experiments, slides were coated with a poly-D-lysine mixture as previously described^7^. Briefly, a mixture of 750 µL of LC-MS grade water, 750 µL of poly-D-lysine and 1 µL of IGEPAL CO-630 detergent was vortexed thoroughly. 20 µL of the prepared mixture was spotted and spread homogenously across each slide using a pipette tip. Samples were dried using a hot plate set to 80°C for approximately 5 minutes. Freshly coated slides were rinsed twice in LC-MS grade water and dried in a speed-vac for 10 minutes. Slides were stored at 4°C until used for cell culture.

For cells grown directly onto slides, silicon 8-well adapters were attached to poly-D-lysine coated ITO slides. To each well, 200 µL of supplemented cell culture media with 10,000 cells were added and allowed to adhere overnight. After adherence, slides were dipped twice in sterile PBS to remove media before fixing in 4 % paraformaldehyde for 10 minutes. After fixation, slides were dipped twice in 50 mM ammonium formate to enhance lipid signal and twice in LC-MS grade water before storing at -80 °C until lipid MSI analysis. For cytospin samples, a total volume of 120 µL of a 1×10^5^ cells per milliliter suspension was loaded into a cytofunnel attached to ITO slides and spun at 600 rpm for 3 minutes. Samples underwent the same wash, and fixation steps as poly-D-lysine samples prior to storage at -80°C until MSI analysis.

### Preparation for Protein Analysis Utilizing MALDI-IHC Probes

For the analysis of proteins, fresh hLF samples grown on poly-D-lysine slides were immediately prepared for MALDI-IHC analysis following AmberGen’s optimized protocol with minor adaptations. Samples were taken from the incubator and washed twice with PBS, fixed for 10 minutes in 10% paraformaldehyde, then washed for 10 minutes in 1x PBS, two 3-minute washes in acetone, and a 3-minute wash in Carnoy’s solution (6:3:1 Ethanol: Chloroform: Acetic Acid) to remove lipid signals. Samples were rehydrated using a series of washing steps in 100% ethanol (2 × 3min), 95% ethanol (1 × 3 min), 70% ethanol (1 × 3 min), 50% ethanol (1 × 3 min), and 1X TBS (2 × 3 min) using a HistoPro 414. Antigen retrieval was performed using Alkaline-EDTA buffer within a polypropylene Coplin staining jar preheated to 95°C for 30 minutes. Then, the Coplin jar was removed from heat and allowed to cool to room temperature for another 30 minutes. After antigen retrieval, slides were washed in 1X TBS for 10 minutes. Moisture was wicked away from the slide using a Kimwipe and a hydrophobic (PAP) barrier pen was utilized to create a barrier surrounding the cell areas. Samples were then treated with 100 µL of tissue blocking buffer (2% (v/v) mouse and rabbit sera, 5% (w/v) BSA in TBS-OBG) for 1 hour. Working probe mixture was prepared at a 2.0 µg/mL final concentration in the tissue blocking buffer and clarified using a spin filter. After blocking, tissue blocking buffer was carefully removed with a pipette and replaced with 150 µL of working probe mixture and incubated in a humid chamber at 4°C for 18 hours.

After an 18-hour incubation period, samples were washed using the autostainer with 1X TBS (3 × 3 minutes) and 50 mM ammonium bicarbonate buffer (3 × 3 minutes). Samples were dried in the dark in a vacuum desiccator for 1 hour and 30 minutes before photocleaving the mass tags using the AmberGen Light Box for 10 minutes.

### Matrix Application for Lipid and Protein Analyses

For lipid analysis, slides were coated with 1,5-Diaminonapthalene (DAN) matrix at a concentration of 10 mg/mL in 90% acetonitrile using an HTX imaging M5 TM-Sprayer (Chapel Hill, NC) or N-(1-napthyl)ethylenediamine dihydrochloride (NEDC) at a concentration of 10 mg/mL in 90% methanol with 100 mM ammonium fluoride. For AmberGen protein analysis, slides were coated with α-cyano-4-hydroxycinnamic acid (CHCA) at a concentration of 10 mg/mL in 70% acetonitrile with 0.1% trifluoroacetic acid. CHCA matrix was recrystallized using 1 mL of 5% isopropanol in an oven at 55°C for 1 minute and 30 seconds. Sprayer parameters can be found in the supporting information.

### Data Collection

MALDI-MSI spectra were acquired using a timsTOF fleX MALDI-2 mass spectrometer (Bruker Daltonics, Bremen, Germany). External calibration was completed using Agilent ESI-TOF calibration standard. Instrument parameters for each experiment can be found in **Supplementary Table 1**. For each sample type, 3 areas of approximately 10,000 pixels were analyzed to represent technical triplicate. Data was analyzed in SCiLS lab 2023b and normalized using root mean square (MALDI-IHC) or matrix clusters (lipids). Data segmentation was completed in SCiLS using weak denoising and bisecting k-means analysis. Putative identifications were determined using the LipidMaps-2017-12-12 database within Metaspace2020^13^.

## RESULTS AND DISCUSSION

### Comparison of Single Cell Preparation Methods

To determine the best methodology for preparing single-cell samples for MSI analysis, a comparison of utilizing poly-D-lysine as opposed to a cytospin preparation was conducted in hLFs. While the cytospin prepared cells faster, the cell architecture of the characteristic fibroblast branching was not observed. As can be observed in **Figure 2** (panels **2A** and **2C**), when prepared using the cytospin, cells are spherical in shape, and maintain the spherical structures observed when cells are enzymatically detached with trypsin. In comparison, cells that are allowed to attach overnight begin to branch outwards while adhering to the slides **(Figure 2B** and **2C**). Out of the 50 most abundant peaks detected during imaging runs, both preparation methods exhibited similar lipid signals and masses that distinguished the branching cytoplasm and cell nucleus (**Supporting Information Table I**). In comparing the signal intensity for lipids within the two preparation methods, masses that localized to the putative cell nucleus were more intense in the cytospin preparation. This increase in intensity may be due to the stretching of the nucleus within branched fibroblasts, lowering the intensity of the signal observed (intensity boxplots for representative masses used in the blended image can be found in supporting information **Figure S2**).

**Figure 2:**
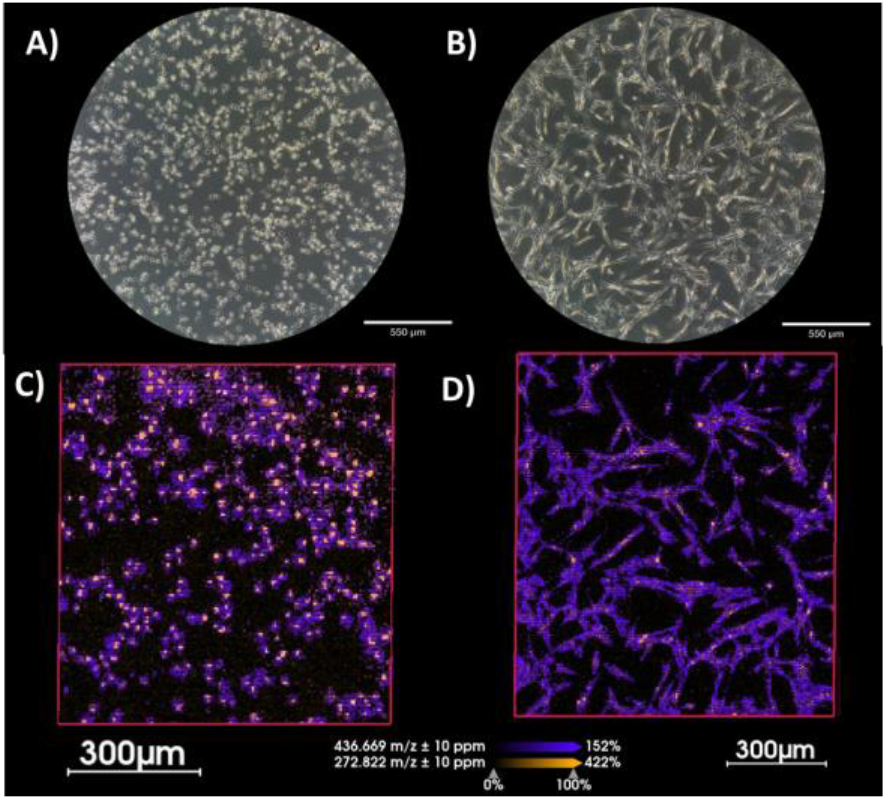
Comparison of hLF cell architecture when prepared using a cytospin (A/C) or when grown onto slides coated with poly-D-lysine (B/D). Images A/B were taken at 10X objective on a brightfield microscope. MALDI-MSI images taken on timsTOF flex with MALDI-2 using DAN matrix at 5 µm resolution of each preparation method and normalized by root mean square. Displayed here are putative lipid species at 272.821 Da ± 10 ppm (orange) localizing primarily to the nucleus and 436.669 Da ± 10 ppm (purple) localizing to cytoplasm.

While the use of both cell preparations for MSI are viable, we selected cells grown onto poly-D-lysine coated slides for the remainder of this study. Though it has a longer protocol, this method was selected based on the branching architecture of the cells. The branching structure provides a larger area for analysis and better mimics the fibroblasts’ native state within lung tissue.

### Analysis of Vimentin and p21 Expression in Fibroblasts

To determine the utility of the MALDI-IHC platform for measuring senescence-related protein levels, we analyzed vimentin and p21 expression in cell lines from three donors representing healthy young, healthy old, and replicative senescent fibroblasts. This approach builds on previous studies that have demonstrated elevated vimentin and p21 levels in senescent fibroblasts compared to younger cells^14^. To ensure that the background signal from the slide did not alter the analysis, segmentation of just the fibroblast species was completed using 12 masses found in the average spectrum (**Figure S3**). As can be seen in **Figure 3**, signal from mass-tagged vimentin slightly increased in the senescent cell lines, with higher incidence of signal coming from the branching cytoplasm. Statistical significance was not observed in the mass spectrometry data. A boxplot of all pixel intensities used in the analysis can be found in the supporting information (**Figure S3-B**). Complementary western blot and immunofluorescence analyses confirmed that the expression of vimentin was not statistically significant across age populations, highlighting the variability among human samples. However, a slight increase in vimentin levels was observed in the replicative senescence group, which may be attributed to low variability within the model. Furthermore, hLF at later passages exhibited elevated vimentin expression as they progressed through senescence (**Figure S4**).

**Figure 3:**
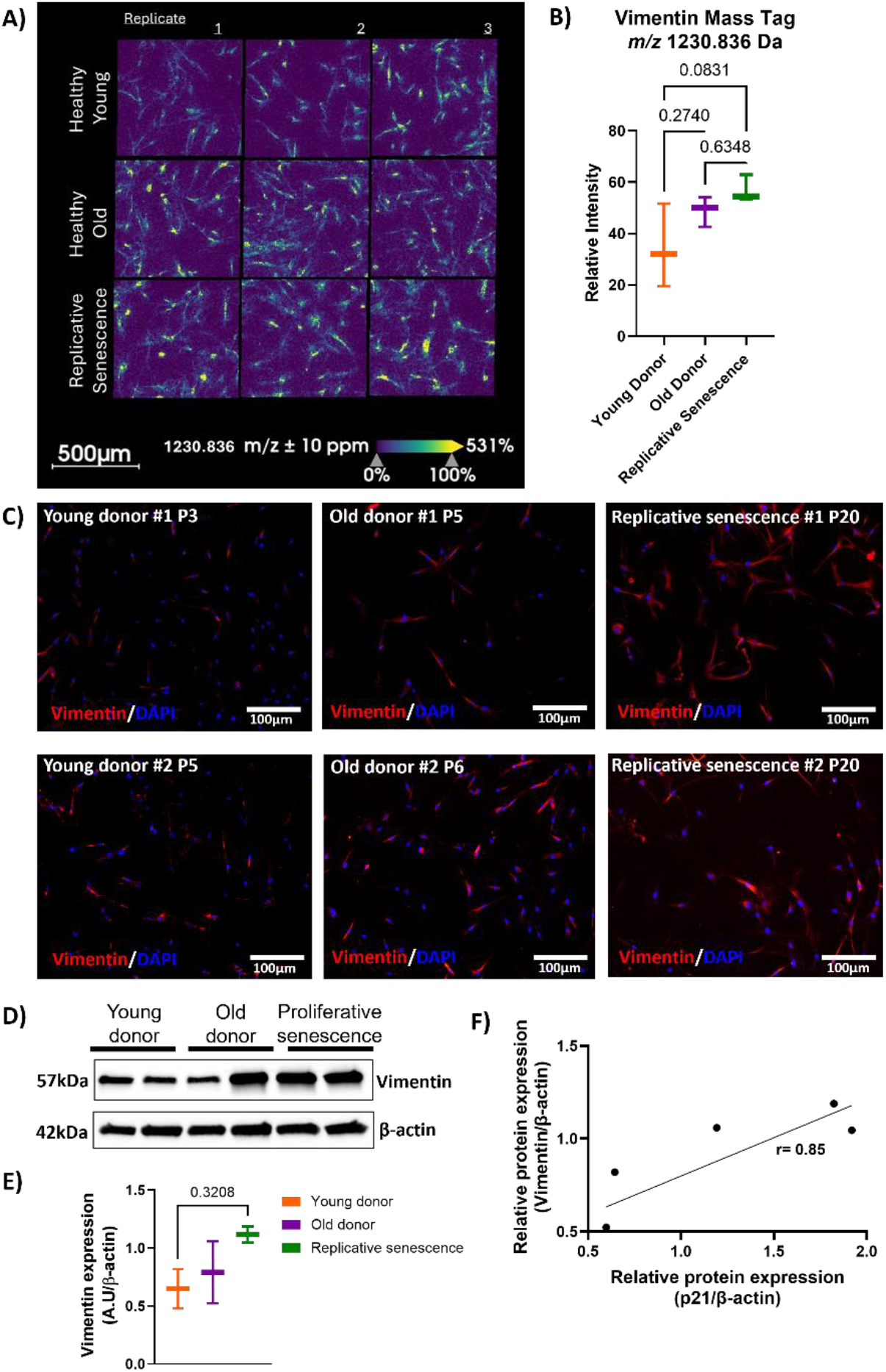
Analyses of vimentin in hLF. **A)** Expression of vimentin as observed through the utilization of MALDI-IHC photo-cleavable mass tag at 1230 Da. **B)** Statistical analysis of spectral intensity from fibroblasts showed vimentin expression was non-significant between groups but a slight increase in average intensity was observed. **C)** Immunofluorescence staining of vimentin in hLF from young and old donors, as well as cells in replicative senescence (x10 magnification, 100µm). Vimentin is shown in red, and nuclei are stained with DAPI in blue. **D)** Western blot analysis of vimentin protein levels in hLF homogenates from all groups. **E)** Quantitative densitometry of vimentin expression, with β-actin used as a loading control. **F)** Correlation between vimentin and p21 expression, with a Pearson correlation coefficient of r=0.85. Data represent mean value ± SEM. Statistical significance was determined by one-way ANOVA followed by post-hoc Tukey, p < 0.05.

Given the Pearson correlation coefficient of 0.85 between vimentin expression and p21 expression, which indicates a positive relationship between these proteins, we decided to incorporate p21—a well-established senescence marker—into our analysis (**Figure 3F**).

Within this group of samples, a matching young donor’s samples for early passage (5) and late passage (20) were utilized for the comparison in the MALDI data. The samples were stained with both vimentin and the p21 antibodies for this analysis. To our knowledge, this is the first application of p21 antibody in MALDI-IHC. The vimentin signal was again not statistically significant between the healthy young and replicative senescent groups. In comparison, the p21 analysis by MALDI-IHC exhibited a statistically significant signal alteration between groups (**Figure 4A/C**). The validation of the p21 antibody was completed on all six of the donors utilizing immunofluorescence and western blot analyses to compare healthy young, healthy old, and replicative senescence populations of fibroblasts. As can be seen in **Figure 5**, the percent of cells positive for p21 was statistically significant between all groups. Furthermore, statistical significance was observed between p21 expression of the healthy young and induced replicative senescence groups. These experiments show the utility of MALDI-IHC probes for relative quantitation in cultured cell systems, specifically within the context of senescence.

**Figure 4:**
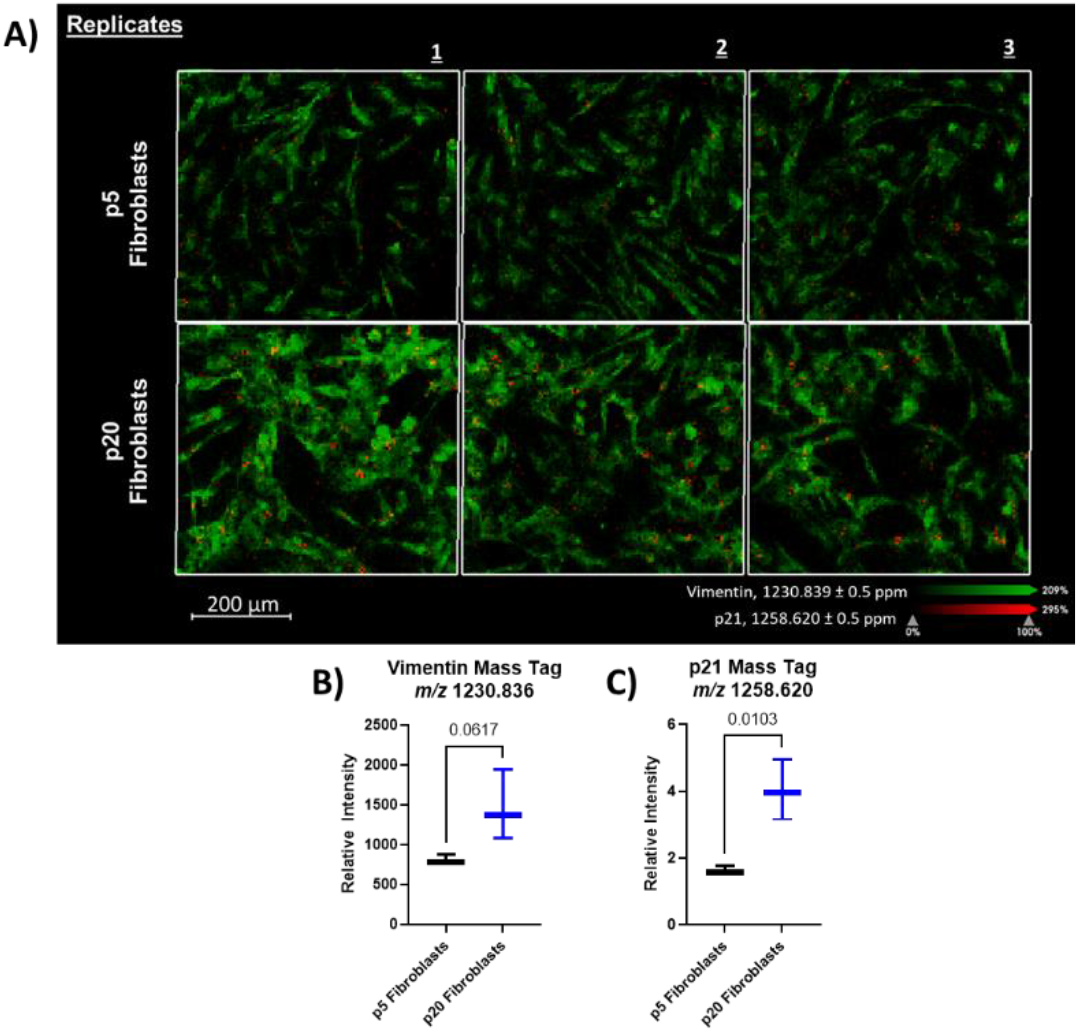
Analysis of p21 expression as a senescence marker in hLF by MALDI-IHC. **A)** MALDI MSI of AmberGen photocleavable mass tags to evaluate the expression of p21, indicated in red at *m/z* 1258.620 Da± 0.5 ppm, and vimentin, indicated in green at *m/z* 1230.839 Da ± 0.5 ppm. MALDI MSI analyses were completed on a timsTOF fleX at 5µm resolution with CHCA matrix, TIC normalized. **B)** Statistical analysis of the observed vimentin mass tag signal intensity for fibroblasts in early and late passages. **C)** Statistical analysis of the observed p21 mass tag signal intensity for fibroblasts in early and late passages.

**Figure 5:**
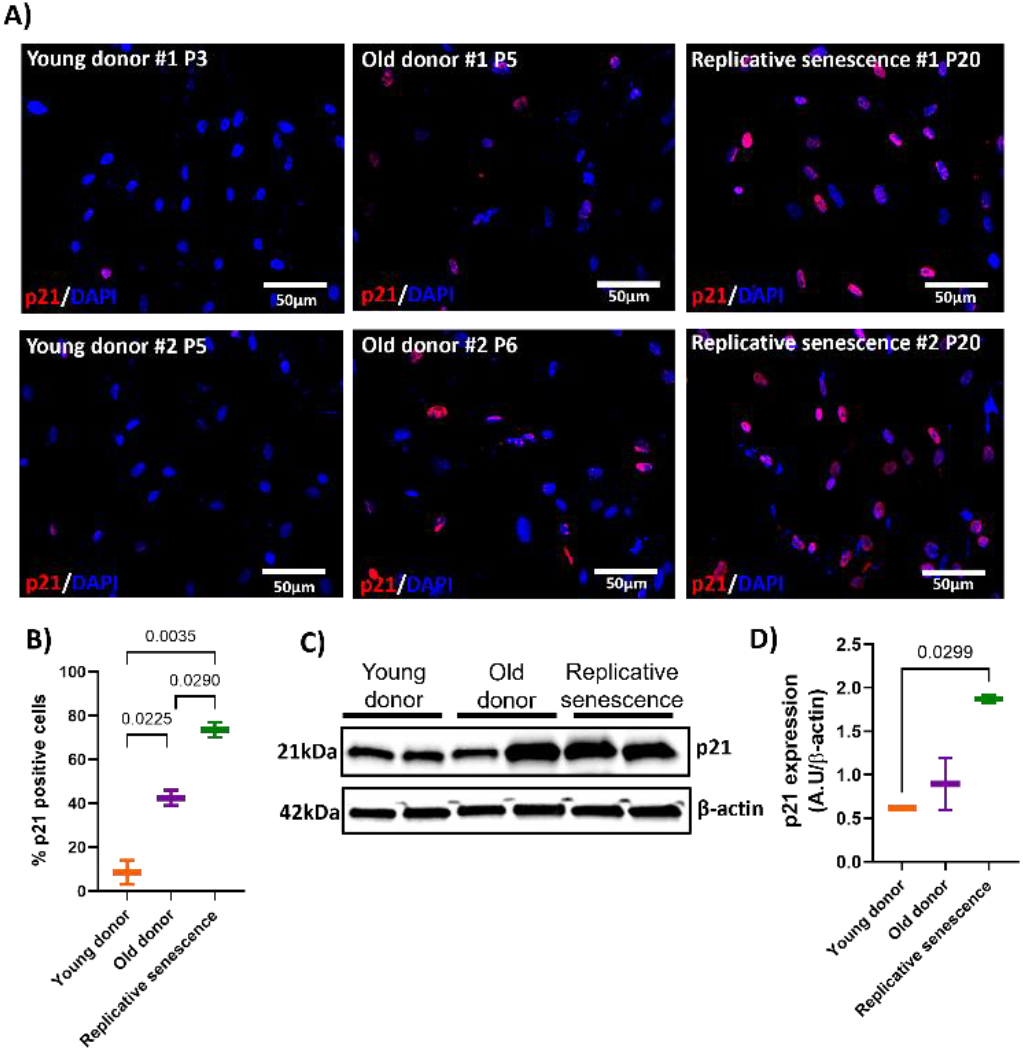
Analysis of p21 expression as a senescence marker in hLF by immunofluorescence and western blot. **A)** Immunofluorescence analysis of p21 in hLF from young and old donors, and cells in replicative senescence. Images were captured at 20x magnification with a scale bar of 50µm. p21 is indicated in red, and nuclei are labeled with DAPI in blue. **B)** Percentage of p21-positive cells in hLF from young and old donors, as well as in cells undergoing replicative senescence. **C)** Western blot analysis of p21 protein levels in hLF homogenates from all groups. **D)** Quantitative densitometry of p21 expression, with β-actin as the loading control. Data represent mean value ± SEM. Statistical significance was determined by one-way ANOVA followed by post-hoc Tukey, p < 0.05.

### Changes in Lipid Signal due to Senescence

Within the observed lipidomic dataset, it was imperative to separate the cell signal from the background to determine changes occurring in cells due to senescence. Normalization techniques were compared (RMS, TIC, and using peaks from matrix clusters^15^) and the utilization of TIC normalization was used throughout the lipid study. Next, by choosing a small set of 20 lipids observed within regions of the cells, unbiased segmentation of data was obtained for subsequent analyses (**Figure S5**). For each replicate, data sets were segmented, and average intensities of lipid signals were exported for further analysis within GraphPad using a student’s t-test (2-tailed)(**Supporting Table S2**). In the initial dataset using DAN as a matrix, very few statistically significant alterations were observed in this mono-culture system, but lipids commonly observed within senescence were detected within the fibroblasts (**Figure 6**)^1, 16^. For example, previous studies examining replicative senescence in dermal fibroblasts observed classes including fatty acids, phosphatidylserines, and phosphatidic acids, which were also detected in our dataset.^16^ Other notable species in this paper correlated to lipids which preferentially ionize in positive ion mode and will be interrogated in future studies.

**Figure 6:**
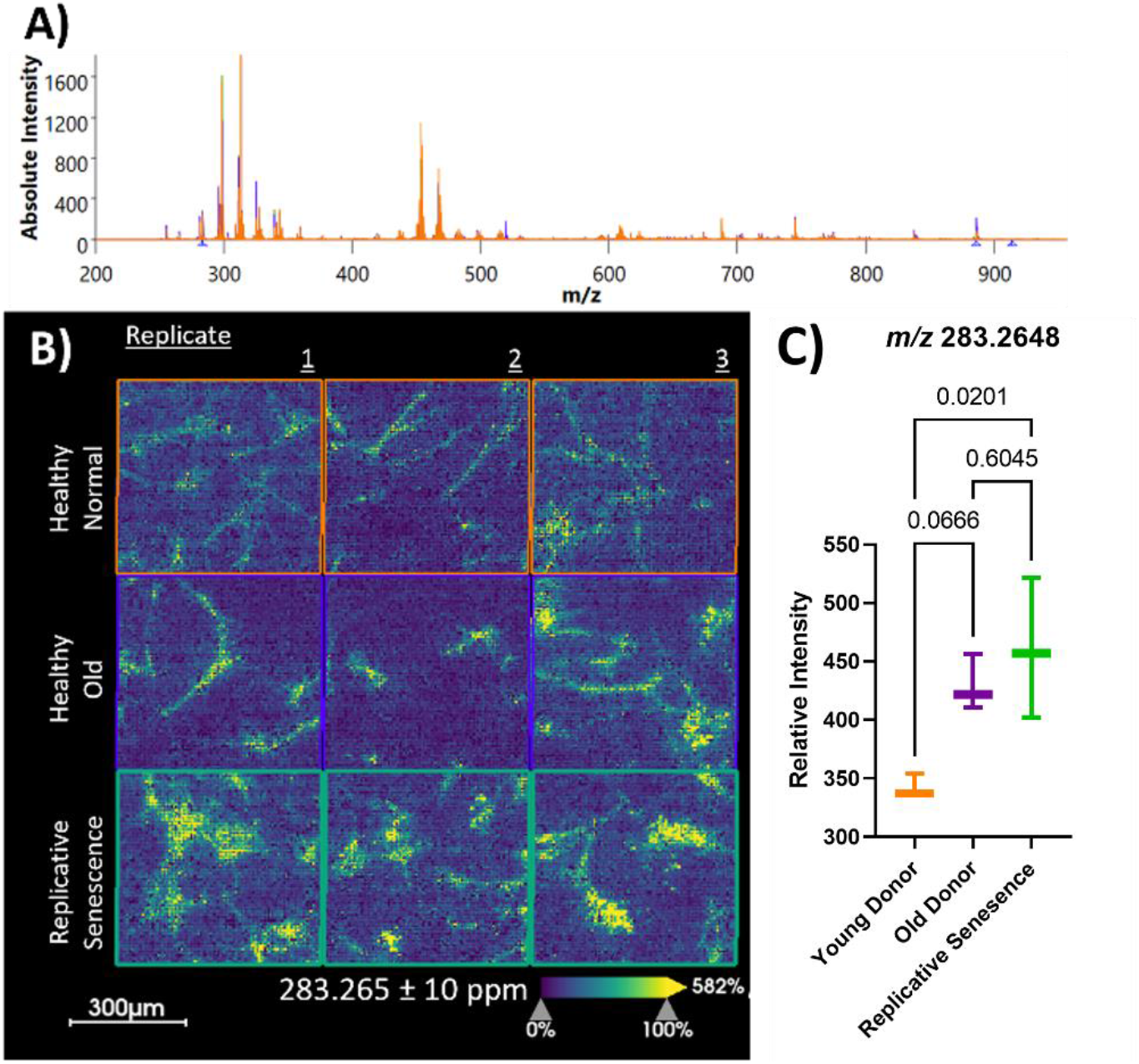
Observation of lipids previously reported within senescence. **A)** Comparison of average mass spectra from the segmented cells for each subtype. Spectra Colors: Orange -healthy young donor, purple - healthy old donor, and green -replicative senescence. **B)** Example of observed signal for putative stearic acid (283.265 Da ± 10 ppm) found to be statistically significant between healthy young donor and replicative senescence fibroblasts **(C)**.

A second analysis was conducted with a different MALDI matrix and the addition of a co-matrix additive, ammonium fluoride, to enhance lipid signals observed from the cell populations. Within this study, of the 44 masses interrogated, 12 were found to be statistically significant (**Supplemental Table S2**). Of the 44 masses detected, 39 were previously reported in senescent cells.^16^ Moreover, ten of these lipid species were found to be in concordance with previous lipidomic analyses of a model of replicative senescence in dermal fibroblasts. A table containing all lipid species tested can be found in **Supporting Table II**.

Due to the high degree of similarity between these cell lines, it is probable that significant alterations to the lipidome may be difficult to detect through a relative label free quantitation study by MSI. Testing of a larger area of cells in tandem with the use of deuterated lipid standards for more stringent quantitation may provide stronger statistical significance between these populations. Furthermore, as lipid levels are controlled through complex mechanisms in multiple cell types, it is possible that single cell imaging analyses of monocultures is unable to fully recapitulate significant alterations to lipid levels that are commonly observed within senescent tissues.

## CONCLUSIONS

Advances to mass spectrometry imaging provides a unique opportunity to interrogate omics-based changes at a single cell level. In this study, for the first time, the utilization of MALDI-IHC, confirmed through western blot and immunofluorescence analyses, demonstrated that the platform can be used to analyze senescent samples at a single cultured cell level. MALDI-IHC may also show future potential in detecting protein markers for specific cells within tissue samples to find cellular localizations in complex samples, such as human biopsies. While we purposefully chose a challenging test set to look for subtle differences in cell lipid composition due to senescence, a few statistically significant lipids were observed within the datasets and further work will aim to more accurately quantitate this molecular subclass. These workflows are robust tools for generating more specific and sensitive signatures of senescence heterogeneity due to cellular populations and will enable more precise mapping of senescence within the lung and other tissues.

## ASSOCIATED CONTENT

## Supporting Information

Supporting information includes additional experimental details and MSI images, descriptions of segmentation analyses, fibroblast characterization through western blot analyses and SA-β-gal assays, and tables detailing the statistical significance determined for interrogated lipid species.

The Supporting Information is available free of charge on the ACS Publications website.

## ACKNOWLEDGMENT

E.R.S, J.H.H, and A.B.H. are supported by NIGMS: R01GM110406 and NIA: RF1AG072760. The OSU campus chemical instrument center (CCIC) which houses the timsTOF fleX is supported by P30 CA016058. T.K., L.R., Q.D.A., M.R., and A.L.M are supported by R01HL149825 and U01HL145550. TOC graphic and workflow created using BioRender.com.

## For table of contents use only

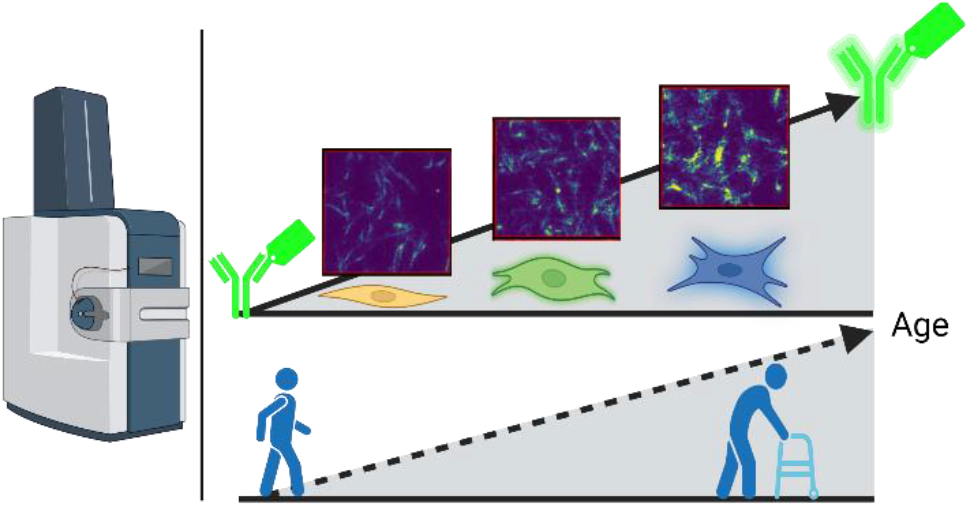

## REFERENCES

(1) Hamsanathan, S.; Gurkar, A. U. Lipids as Regulators of Cellular Senescence. Front Physiol 2022, 13, 796850. DOI: 10.3389/fphys.2022.796850.

(2) Hayflick, L.; Moorhead, P. S. The serial cultivation of human diploid cell strains. Exp Cell Res 1961, 25, 585–621. DOI: 10.1016/0014-4827(61)90192-6.

(3) Suryadevara, V.; Hudgins, A. D.; Rajesh, A.; Pappalardo, A.; Karpova, A.; Dey, A. K.; Hertzel, A.; Agudelo, A.; Rocha, A.; Soygur, B.; et al. SenNet recommendations for detecting senescent cells in different tissues. Nat Rev Mol Cell Biol 2024. DOI: 10.1038/s41580-024-00738-8.

(4) Sliogeryte, K.; Gavara, N. Vimentin Plays a Crucial Role in Fibroblast Ageing by Regulating Biophysical Properties and Cell Migration. Cells 2019, 8 (10). DOI: 10.3390/cells8101164.

(5) Nishio, K.; Inoue, A.; Qiao, S.; Kondo, H.; Mimura, A. Senescence and cytoskeleton: overproduction of vimentin induces senescent-like morphology in human fibroblasts. Histochem Cell Biol 2001, 116 (4), 321–327. DOI: 10.1007/s004180100325.

(6) Taylor, M. J.; Lukowski, J. K.; Anderton, C. R. Spatially Resolved Mass Spectrometry at the Single Cell: Recent Innovations in Proteomics and Metabolomics. J Am Soc Mass Spectrom 2021, 32 (4), 872–894. DOI: 10.1021/jasms.0c00439.

(7) Cuypers, E.; Claes, B. S. R.; Biemans, R.; Lieuwes, N. G.; Glunde, K.; Dubois, L.; Heeren, R. M. A. ‘On the Spot’ Digital Pathology of Breast Cancer Based on Single-Cell Mass Spectrometry Imaging. Anal Chem 2022, 94 (16), 6180–6190. DOI: 10.1021/acs.analchem.1c05238.

(8) Capolupo, L.; Khven, I.; Lederer, A. R.; Mazzeo, L.; Glousker, G.; Ho, S.; Russo, F.; Montoya, J. P.; Bhandari, D. R.; Bowman, A. P.; et al. Sphingolipids control dermal fibroblast heterogeneity. Science 2022, 376 (6590), eabh1623. DOI: 10.1126/science.abh1623.

(9) Yagnik, G.; Liu, Z.; Rothschild, K. J.; Lim, M. J. Highly Multiplexed Immunohistochemical MALDI-MS Imaging of Biomarkers in Tissues. J Am Soc Mass Spectrom 2021, 32 (4), 977–988. DOI: 10.1021/jasms.0c00473.

(10) Claes, B. S. R.; Krestensen, K. K.; Yagnik, G.; Grgic, A.; Kuik, C.; Lim, M. J.; Rothschild, K. J.; Vandenbosch, M.; Heeren, R. M. A. MALDI-IHC-Guided In-Depth Spatial Proteomics: Targeted and Untargeted MSI Combined. Anal Chem 2023, 95 (4), 2329–2338. DOI: 10.1021/acs.analchem.2c04220.

(11) Lim, M. J.; Yagnik, G.; Henkel, C.; Frost, S. F.; Bien, T.; Rothschild, K. J. MALDI HiPLEX-IHC: multiomic and multimodal imaging of targeted intact proteins in tissues. Front Chem 2023, 11, 1182404. DOI: 10.3389/fchem.2023.1182404.

(12) Dunne, J.; Griner, J.; Romeo, M.; Macdonald, J.; Krieg, C.; Lim, M.; Yagnik, G.; Rothschild, K. J.; Drake, R. R.; Mehta, A. S.; et al. Evaluation of antibody-based single cell type imaging techniques coupled to multiplexed imaging of N-glycans and collagen peptides by matrix-assisted laser desorption/ionization mass spectrometry imaging. Anal Bioanal Chem 2023, 415 (28), 7011–7024. DOI: 10.1007/s00216-023-04983-2.

(13) Palmer, A.; Phapale, P.; Chernyavsky, I.; Lavigne, R.; Fay, D.; Tarasov, A.; Kovalev, V.; Fuchser, J.; Nikolenko, S.; Pineau, C.; et al. FDR-controlled metabolite annotation for high-resolution imaging mass spectrometry. Nat Methods 2017, 14 (1), 57–60. DOI: 10.1038/nmeth.4072.

(14) Stein, G. H.; Drullinger, L. F.; Soulard, A.; Dulic, V. Differential roles for cyclin-dependent kinase inhibitors p21 and p16 in the mechanisms of senescence and differentiation in human fibroblasts. Mol Cell Biol 1999, 19 (3), 2109–2117. DOI: 10.1128/MCB.19.3.2109.

(15) Treu, A.; Rompp, A. Matrix ions as internal standard for high mass accuracy matrix-assisted laser desorption/ionization mass spectrometry imaging. Rapid Commun Mass Spectrom 2021, 35 (16), e9110. DOI: 10.1002/rcm.9110.

(16) Lizardo, D. Y.; Lin, Y. L.; Gokcumen, O.; Atilla-Gokcumen, G. E. Regulation of lipids is central to replicative senescence. Mol Biosyst 2017, 13 (3), 498–509. DOI: 10.1039/c6mb00842a.

