## Supporting Information for "Single Cell MALDI-MSI Analysis of Lipids and Proteins within a Replicative Senescence Fibroblast Model"

**Reagents and Materials**:

ITO slides were obtained from Delta Technologies Ltd. (Loveland, CO, USA). HPLC grade water, methanol, and acetonitrile were obtained from Burdick & Jackson (Honeywell Research Chemicals, Muskegon, MI, USA). Poly-D-Lysine, phosphate-buffered saline (PBS), 0.05% trypsin, fetal bovine serum, F-12 nutrient Ham medium, and antimycotic-antibiotic were obtained from Gibco (Waltham, MA, USA). Dimethyl sulfoxide (DMSO), IGEPAL CA-630, powder paraformaldehyde, ammonium formate, reagent grade ethanol, acetic acid, 99.99%+, concentrated alkaline retrieval buffer, hydrophobic (PAP) barrier pen, bovine serum albumin (BSA), 50% octyl β-D-glucopyranoside solution (OBG), 1,5-Diaminonapthalene (DAN), α-cyano-4-hydroxycinnamic acid (CHCA),N-(1-napthyl)etylenediamine dihydrochloride (NEDC), ammonium fluoride, sodium chloride >99.5%, Triton X-100, β-actin (A3854), and ammonium bicarbonate were obtained from Sigma-Aldrich (St. Louis, MO, USA). HPLC Grade chloroform, HPLC and GC grade acetone were obtained from Fisher Chemical (Hampton, NH, USA). Normal rabbit serum and normal mouse serum were obtained from Jackson ImmunoResearch Laboratories Inc. (West Grove, PA, USA). Molecular biology grade tris-HCL and Molecular tris base were obtained from Promega (Madison, WI, USA). MALDI-IHC probes were obtained from AmberGen (Billerica, MA, USA). Senescence β-galactosidase staining kit (#9860), vimentin antibody (5741), p21 antibody (2947), and horseradish peroxide-conjugated secondary antibody (7074) were obtained from Cell Signaling Technology (Danvers, MA, USA). RIPA Buffer Lysis System (sc-24948) was obtained from Santa Cruz Biotechnology, Inc. (Dallas, TX, USA).Pierce™ BCA Protein Assay (#23228) was obtained from Thermo Fisher Scientific (Waltham, MA, USA). Non-fat dry milk (Dry Powder Milk) was obtained from Research Products International (Mount Prospect, IL, USA). Alexa Fluor 647-conjugated secondary antibody, SlowFade™ Gold Antifade, 4',6-Diamidino-2-Phenylindole, Dihydrochloride (DAPI), and EVOS™ microscope were obtained from Invitrogen (Carlsbad, CA, USA). Nitrocellulose membrane, Trans-Blot Turbo™ transfer system, and Clarity™ Western ECL substrate (1705061) were obtained from Bio-Rad (Hercules, CA, USA).

**Cell Culture Growth and Treatments**

After tissue collection, enzymatic digestion (0.05% trypsin) was used to isolate primary human lung fibroblast (hLFs), from young (<30 years old) and old (>60 years old) donors. Replicative senescence was induced by passaging the sample obtained from a young donor. Early (passage 3-5) and late (passage 20) passage cultures of hLFs were grown with F-12 nutrient Ham Medium containing 10% of fetal bovine serum, and 1% of antimycotic-antibiotic. Cells were cultured at 37 °C, 5% $CO_2$ and then expanded to select a homogenous fibroblast population. To detect cellular senescence, cells were stained with SA-β gal according to the manufacturer's instructions for the senescence β-galactosidase staining kit. Images were acquired with color brightfield on an EVOS™ microscope (M7000) at 10X magnification. The percentage of SA-β gal positive cells was quantified using Image J software (Bethesda, MD) (Figure S1).

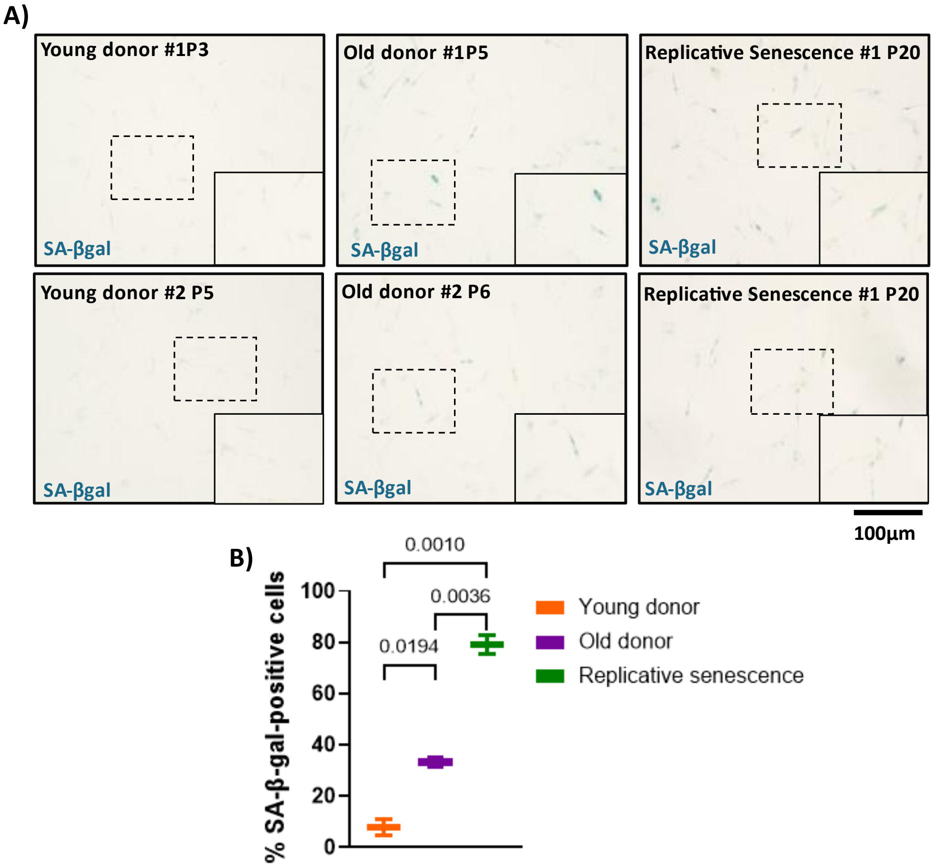


**Figure S1: Assessment of β-galactosidase activity as a marker of cellular senescence in hLF. A)** SA-βgal staining in hLF from young and old donors, as well as in cells undergoing replicative senescence. Images were captured at 10x magnification with a scale bar of 100µm. **B)** Percentage of SA-β-gal-positive cells in hLF from all groups. Data represent mean value ± SEM. Statistical significance was determined by one-way ANOVA followed by post-hoc Tukey, $p < 0.05$.

**Matrix Application – Sprayer Notes**

For DAN, matrix was applied at 70°C for 2 passes over the samples. The flow rate of the matrix was 0.12 mL/min at a velocity of 800 mm/min, track spacing of 2 mm, pressure of 10 psi, gas flow rate of 3 L/min, and nozzle height of 40 mm. For CHCA, matrix was applied at 60°C for 8 passes over the samples. The flow rate of the matrix was 0.1 mL/min at a velocity of 1,350 mm/min, track spacing of 3 mm, pressure of 10 psi, gas flow rate of 2 L/min, drying time of 10 sec and nozzle height of 40 mm. For NEDC, matrix was applied at 30°C for 14 passes over the samples. The flow rate of the matrix was 0.06 mL/min at a velocity of 1,200 mm/min, track spacing of 3mm, pressure of 10 psi, gas flow rate of 2 L/min, no drying time was utilized and nozzle height of 40 mm.

**Immunofluorescence**

Fibroblasts were grown on poly-D-lysine coated slides, fixed with 4% paraformaldehyde for 10 minutes, permeabilized with 0.1 % Triton X-100 in PBS (PBS-T) for 15 min and then blocked with 20% donkey serum in 0.5% BSA/PBS for 1 hour at room temperature (RT), followed by incubation overnight with primary antibody against vimentin (1:200) and p21 (1:100) at 4°C. Slides were then stained with Alexa Fluor 647-conjugated secondary antibody (1:250). For identification of nuclei, DAPI (1:2000) was applied for 10 min. Coverslips were applied to slides using SlowFade™ Gold Antifade, and tissues were visualized using an EVOS$^{TM}$ microscope (M7000). Images were acquired using a 10X objective and quantified using the software Image J (NIH, Bethesda, MD).

**Western blot**

Fibroblasts were grown on a T75 Flask until 80% confluence, and cells were washed three times with PBS and 100μL of RIPA buffer containing 1% of protease and phosphatase inhibitors. After scraping, the lysate was collected and frozen. On the day the assay was performed, protein concentration was quantified by Pierce™ BCA Protein Assay. Equal amounts of protein from cell preparations were separated by sodium dodecyl sulfate-polyacrylamide gel electrophoresis (SDS-PAGE) and electrotransferred to a nitrocellulose membrane using a Trans-Blot Turbo™ transfer system. After transfer, membranes were washed in TBS-T (10 mM Tris, pH 8.0, 150 mM NaCl, 0.05 % Tween 20) and blocked with TBS-T supplemented with 5 % non-fat dry milk for 1 hour at RT. Subsequently, the membranes were incubated overnight at 4 °C with primary antibodies in TBS-T against vimentin (1:1000), p21 (1:1000), and β-actin (1:30,000). The next day, the membranes were washed and incubated with horseradish peroxide-conjugated secondary antibody (1:2000) for 1 hour at RT. Following additional wash steps with TBS-T, membranes were treated with Clarity™ Western ECL substrate, and chemiluminescence was detected (Chemidoc System, Bio-Rad). Quantification was performed by signal intensity measurement with Image J software (NIH, Bethesda, MD)

**Statistical Analysis for Immunofluorescence and Western Blot Analyses**

Results are expressed as a mean ± standard error of the mean (SEM). Differences between means of the three groups were examined using a one-way analysis of variance (ANOVA) followed by the Tukey post hoc test. Statistical significance was set at $p<0.05$. Statistical and Pearson correlation analyses were conducted with GraphPad Prism 10.2.0 (GraphPad Software, Inc., San Diego, CA).

| **Application** | MALDI-IHC | Lipid Imaging (DAN) | Lipid Imaging (NEDC) |
|---|---|---|---|
| **Polarity** | Positive | Negative | Negative |
| ***m/z* Range** | 1000 – 1850 | 100 – 1500 | 200 -1500 |
| **Laser Power** | 65% | 55% | 55% |
| **CID** | N/A | N/A | N/A |
| **Frequency** | 5000 Hz | 1000 Hz | 10,000 Hz |
| **Shots** | 300 | 500 | 250 |
| **Laser Setting** | 5 µm | 5 µm | 5 µm |
| **Raster Setting** | 5 µm | 5 µm | 5 µm |
| **MALDI/ MALDI-2** | MALDI | MALDI-2 | MALDI |

**Table S1:** Instrument parameters used for MALDI experiments on the Bruker timsTOF fleX with MALDI-2.

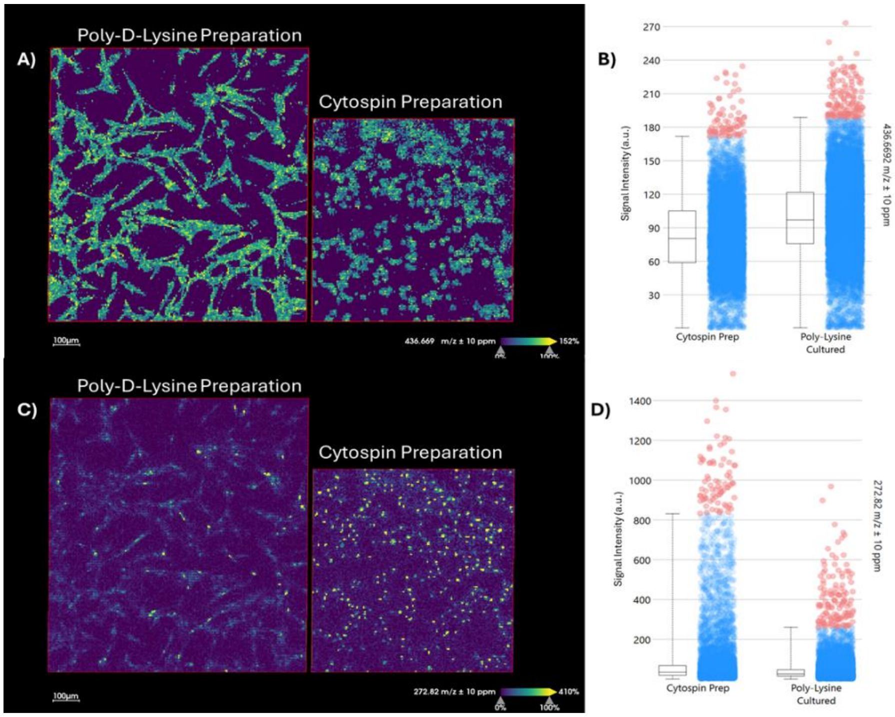


**Figure S2:** Representative mass images of signals correlating to the cytoplasm, 436.669 Da **(A)** and putative nucleus, 272.822 Da **(C)** of cells. Intensity boxplots showing the average signals for the mass images of 436.669 Da **(B)** and 272.822 **(D)**.

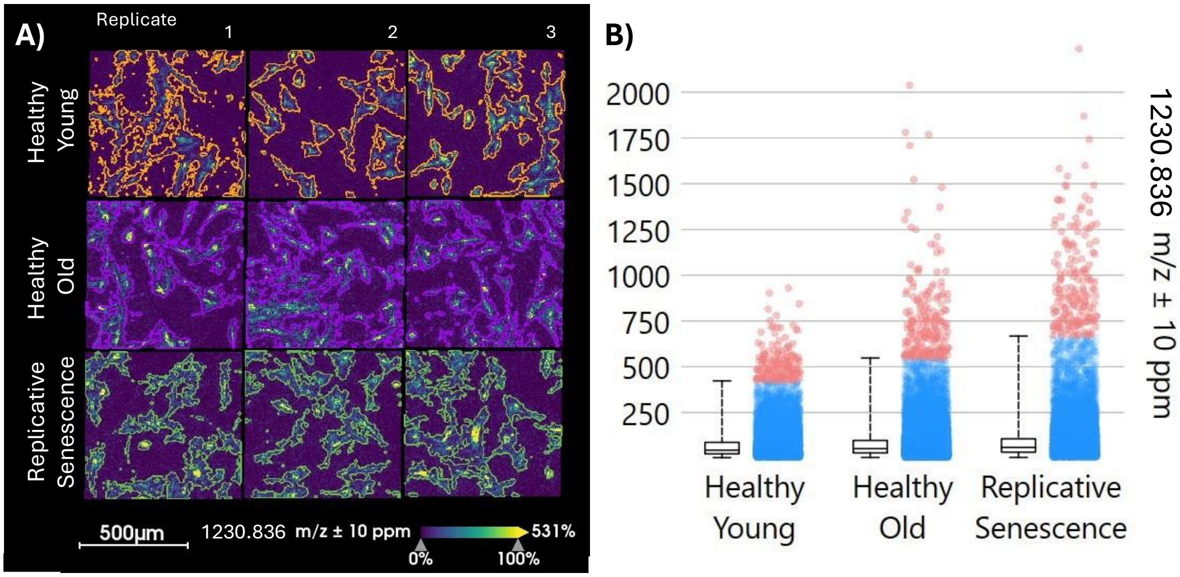


**Figure S3:** Segmentation of fibroblast data in AmberGen analysis **(A)** to find mean spectrum peak intensities to utilize in ANOVA analyses. **B)** Box and whisker plot showing the distribution of intensities for *m/z* 1230.836 Da across three replicate runs of single donors.

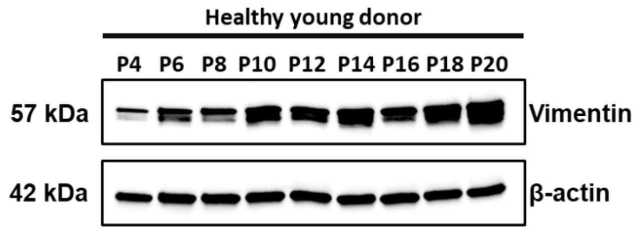


**Figure S4: Vimentin expression undergoing senescence in hLF.** Western blot of vimentin to compare expression levels as a function of passage number with β-actin used as a loading control.

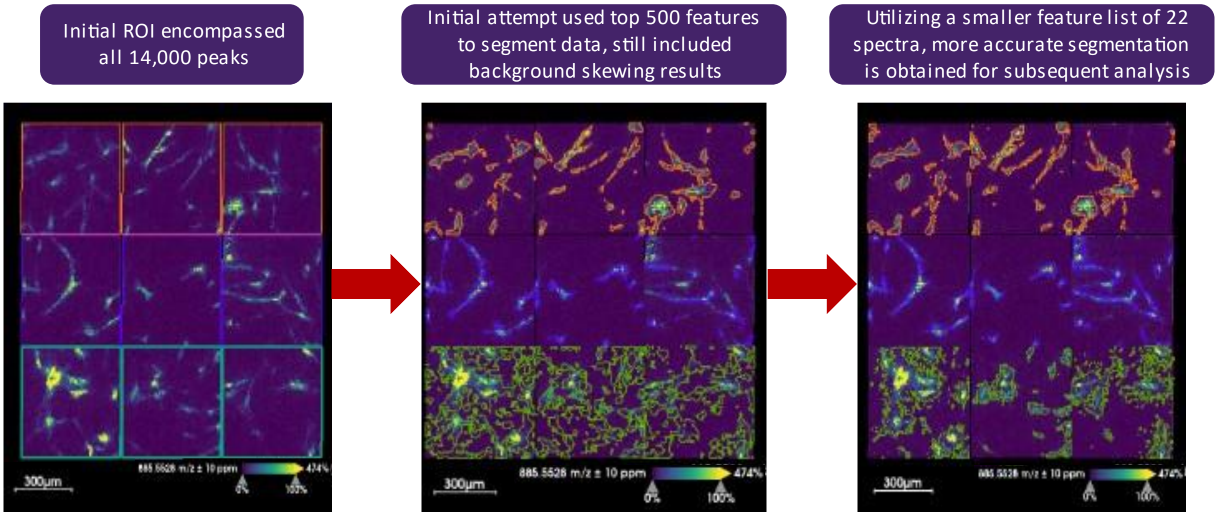


**Figure S5**: Example of data segmentation utilized to ensure that only cell populations were chosen for subsequent statistical analyses. Data normalized utilizing matrix clusters from DAN.

| Observed Mass | ANOVA p-value | Observed Mass | ANOVA p-value |
|---|---|---|---|
| 281.250 | 0.194 | 715.576 | 0.349 |
| 283.265* | 0.021 | 726.581 | 0.177 |
| 303.235 | 0.088 | 728.525 | 0.971 |
| 391.226 | 0.384 | 744.555 | 0.388 |
| 599.320 | 0.088 | 770.571 | 0.692 |
| 616.472 | 0.663 | 772.586 | 0.952 |
| 673.483 | 0.812 | 794.571 | 0.899 |
| 673.526 | 0.443 | 810.683 | 0.099 |
| 687.545 | 0.578 | 857.519 | 0.357 |
| 701.515 | 0.720 | 885.550 | 0.795 |
| 701.561 | 0.342 | 913.581 | 0.513 |

**Table S2**: Statistical assessment of 22 putative lipid species of interest found in MALDI experiments utilizing DAN matrix by one-way ANOVA multiple comparison testing. Mass denoted with * was found to be statistically significant at $p \leq 0.5$.
